# Metagenomic Next-generation Sequencing of Cerebrospinal Fluid for the Diagnosis of Central Nervous System Infections: A Multicentre Prospective Study

**DOI:** 10.1101/658047

**Authors:** Siyuan Fan, Xiaojuan Wang, Yafang Hu, Jingping Shi, Yueli Zou, Weili Zhao, Xiaodong Qiao, Chunjuan Wang, Jerome H. Chin, Lei Liu, Lingzhi Qin, Shengnan Wang, Hongfang Li, Wei Yue, Weihe Zhang, Xiaohua Li, Ying Ge, Honglong Wu, Weijun Chen, Yongjun Li, Tianjia Guan, Shiying Li, Yihan Wu, Gaoya Zhou, Zheng Liu, Yushun Piao, Jianzhao Zhang, Changhong Ren, Li Cui, Caiyun Liu, Haitao Ren, Yanhuan Zhao, Shuo Feng, Haishan Jiang, Jiawei Wang, Hui Bu, Shougang Guo, Bin Peng, Liying Cui, Wei Li, Hongzhi Guan

## Abstract

**Background:** Infectious encephalitis and meningitis are often treated empirically without identification of the causative pathogen. Metagenomic next-generation sequencing (mNGS) is a high throughput technology that enables the detection of pathogens independent of prior clinical or laboratory information.

**Methods:** The present study was a multicentre prospective evaluation of mNGS of cerebrospinal fluid (CSF) for the diagnosis of suspected central nervous system infections.

**Results:** A total of 276 patients were enrolled in this study between Jan 1, 2017 and Jan 1, 2018. Identification of an etiologic pathogen in CSF by mNGS was achieved in 101 patients (36.6%). mNGS detected 11 bacterial species, 7 viral species, 2 fungal species, and 2 parasitic species. The five leading positive detections were varicella-zoster virus (17), *Mycobacterium tuberculosis* (14), herpes simplex virus 1 (12), Epstein-Barr virus (12), and *Cryptococcus neoformans* (7). False positives occurred in 12 (4.3%) patients with bacterial infections known to be widespread in hospital environments. False negatives occurred in 16 (5.8%) patients and included bacterial, viral and fungal aetiologies.

**Conclusions:** mNGS of CSF is a powerful diagnostic method to identify the pathogen for many central nervous system infections.

## 1. INTRODUCTION

Infectious encephalitis and meningitis are major contributors to the neurological global burden of disease^1–4^. Numerous microorganisms, including bacteria, viruses, fungi, and parasites, can cause encephalitis and meningitis in immunocompetent or immunocompromised hosts; but the clinical manifestations of many infections are non-specific. Using comprehensive conventional diagnostic technologies, microbiological detection of the pathogen is achieved in only 50-80% of cases^5–8^. The inability to identify the infectious aetiology of encephalitis and meningitis often results in delayed, inadequate, and/or inappropriate treatment.

Metagenomic next-generation sequencing (mNGS) is a novel tool that allows for the simultaneous and independent sequencing of thousands to billions of DNA fragments^9^. Cerebrospinal fluid (CSF) is particularly suitable for NGS due to its sterility in healthy individuals. Compared with traditional individual target-specific tests, mNGS can identify pathogens without the input of clinical predictors or prior laboratory results. Several recent studies have demonstrated the capability of mNGS of CSF to identify known and unsuspected pathogens and to discover new microorganisms^10–18^. mNGS of CSF is being increasingly utilized in routine clinical settings for the rapid diagnosis of central nervous system infections. However, most published studies are retrospective case reports or case series^11–17,19–24^, and thus, large prospective studies are needed to demonstrate the clinical impact and cost-effectiveness of mNGS for the diagnosis of meningitis and encephalitis. We undertook a multicentre prospective study to comprehensively evaluate the performance of mNGS of CSF for the diagnosis of central nervous system (CNS) infections compared to conventional microbiological methodologies.

## 2. METHODS

### 2.1 Participants and study design

This study was a multicentre prospective cohort assessment of the mNGS of CSF for the diagnosis of suspected infectious encephalitis or meningitis. The participating sites were 20 hospitals located in 10 provinces/municipalities in China. Each hospital is a member of the Beijing Encephalitis Group. Adult patients were eligible for inclusion in the study if they presented with clinical manifestations consistent with either encephalitis or meningitis (Table 1) and if standard diagnostic examinations (Supplementary Table 1) failed to identify an etiological cause within 3 days. Exclusion criteria are shown in Table 1.

**Table 1.** Case definitions and exclusion criteria for encephalitis and meningitis

|  |
| --- |
| **Case definitions for encephalitis** |
| **Major criteria (required)** |
| Patients presenting to medical attention with altered mental status (decreased or altered level of consciousness or personality change) lasting $\geq 24$ hrs with no alternative cause identified, and/or generalized or partial seizures not fully attributable to a pre-existing seizure disorder or a simple febrile seizure. |
| **Minor criteria ($\geq 2$ points)** |
| Documented fever $\geq 38^{\circ}\text{C}$ within the 72 hrs before or after presentation |
| New onset of focal neurologic findings |
| CSF WBC count $\geq 5/\text{cubic mm}$ |
| Abnormality of brain parenchyma on neuroimaging suggestive of encephalitis that is either new from prior studies or appears acute in onset |
| Abnormality on EEG that is consistent with encephalitis and not attributable to another cause. |
| **Exclusion criteria for encephalitis** |
| $\leq 28$ days of age |
| Non-infectious encephalitis, such as autoimmune disorders, paraneoplastic syndromes, NMOSD, neuropsychiatric involvement of rheumatic diseases |
| HIV or syphilis infection |
| History of recent (within 4 weeks before the onset of disease) vaccination |
| Meningitis without clinical brain parenchyma involvement |
| Absolute contraindications for lumbar puncture; |
| Traumatic LP with obvious blood-contaminated CSF |
| Pregnancy |
| Refusal to sign the informed consent |
| **Case definitions for meningitis** |
| Patients presenting to medical attention with at least two of the four symptoms of headache, fever (documented fever $\geq 38^{\circ}\text{C}$ within the 72 hrs before or after presentation), neck stiffness, decreased level of consciousness (defined by a Glasgow Coma Scale score below 14) |
| CSF white blood cell count $\geq 5/\text{cubic mm}$ |
| **Exclusion criteria for meningitis** |
| $\leq 28$ days of age |
| HIV or syphilis infection |
| Meningeal malignancy confirmed by CSF cytology |
| Traumatic LP with obvious blood-contaminated CSF |
| Pregnancy |
| Refusal to sign the informed consent |

mNGS were conducted on all CSF specimens. Relevant conventional microbiological studies (e.g. staining, culture, polymerase chain reaction [PCR], serology) were arranged according to the clinical manifestations and the results of mNGS. Conventional microbiological studies were considered the gold standard according to relevant guidelines and/or consensus^2,25–27^ to classify the results of mNGS as true-positive, false-positive, and false-negative. Detected pathogens were classified as etiologic pathogens if the major clinical manifestations of the patient were consistent with that pathogen. All patients were treated based on the results of conventional microbiological testing (or empirically if results were negative) according to the latest clinical guidelines and/or consensus. Patients were followed for at least 30 days to determine the final diagnosis. Demographic data, medical history, laboratory test results (including all conventional microbiological tests), neuroimaging findings, medical therapy, and response to treatment were collected prospectively. Patients enrolled from Jan 1, 2017 to Jan 1, 2018 were included in the final analyses.

This study was approved by the institutional review board of Peking Union Medical College Hospital (no. JS-890). Written informed consent was obtained from each patient or their legal surrogate prior to enrolment.

### 2.2 mNGS of CSF

CSF samples were collected according to standard sterile procedures, snap-frozen, and stored at −20°C until they were delivered to the sequencing centre. Because reverse transcription was not performed to prepare DNA libraries, RNA viruses were not investigated in this study. mNGS of the CSF samples was performed using a standard flow that has been successfully used to detect herpes simplex virus 1 (HSV1), HSV2, varicella zoster virus (VZV), *Listeria monocytogenes*, *Brucella*, and *Taenia solium*^12–15^.

DNA was extracted from 300 μL of CSF and negative ‘no-template’ controls (NTCs). Sequencing was performed on the BGISEQ-100 platform with an average of 20 million total reads obtained for each sample. The qualified reads were mapped to the human reference genome using the Burrows–Wheeler Aligner to remove human sequences. The remaining reads were aligned to the database of annotation, which includes the NCBI microbial genome database (ftp://ftp.ncbi.nlm.nih.gov/genomes/) to detect pathogens. The sequencing data was analysed in terms of the numbers of raw reads, non-human reads, and reads aligned to the microbial genome database as well as species-specific reads (genus-specific reads for *Mycobacterium tuberculosis* and *Brucella*), reads per million (RPM), and genome coverage (%). The results of mNGS were available in less than 48 hrs.

### 2.3 Criteria for positive results of mNGS of CSF samples

To reduce the influence of potential contamination, we used the following criteria for positive results of CSF mNGS:

1. For extracellular bacteria, fungi (excluding *Cryptococcus*), and parasites, the result was considered positive if a species detected by mNGS had a species-specific read number (SSRN) ≥ 30 (RPM ≥ 1.50) that ranked among the top 10 for bacteria, fungi, or parasites. Organisms detected in the NTC or that were present in ≥ 25% of samples from the previous 30 days were excluded but only if the detected SSRN was ≥ 10-fold than that in the NTC^28^ or other organisms. Additionally, organisms present in ≥ 75% of samples from the previous 30 days were excluded.
2. For intracellular bacteria (excluding *Mycobacterium tuberculosis* and *Brucella*) and *Cryptococcus*, the result was considered positive if a species detected by NGS had a SSRN ≥ 10 (RPM ≥ 0.50)^13^ that ranked among the top 10 for bacteria or fungi. Pathogens detected in the NTC or that were present in ≥ 25% of samples from the previous 30 days were excluded but only if the detected SSRN was ≥ 10-fold than that in the NTC or other organisms.
3. For viruses, *Brucella*, and *Mycobacterium tuberculosis*, the result was considered positive if a species (or genus for *Mycobacterium tuberculosis* [*Mycobacterium tuberculosis* complex, MTC] and *Brucella*) detected by NGS had a SSRN ≥ 3 (RPM ≥ 0.15)^12,28^. Pathogens detected in the NTC were excluded but only if the detected SSRN was ≥ 10-fold than that in the NTC. In our previous clinical observations, there were a few cases without *Mycobacterium tuberculosis* infection which contained MTC-specific reads number of 1 in the mNGS results. To mitigate the possibility of false positives, we adopted the criteria of SSRN ≥ 3 rather than SSRN ≥ 1^24^ in this study. The performance of the criteria were evaluated at the finally stage of the study, the original results of mNGS and/or clinical manifestations were used to guide the further testing of conventional microbiological studies.

### 2.4 Statistical analysis

All statistical analyses were conducted using Statistical Package for the Social Sciences (SPSS) version 17.0 and EXCEL 1810. Depending on their distribution, all data are expressed as medians with interquartile ranges (IQRs) or as means ± standard deviation.

## 3. RESULTS

### 3.1 Characteristics of the study participants

287 patients were screened for inclusion in this study (Fig. 1). 11 patients were initially thought to have CNS infections and mNGS was performed. However, these cases were ultimately excluded following the final diagnosis of a non-infectious disease. Of these 11 excluded patients, 10 had negative mNGS results, and 1 patient receiving immunosuppressive therapy was positive for BK polyomavirus. The final cohort included 276 patients in the study. 176 (63.8%) were male and the median age was 42 years (IQR: 26–54 years). The median time from disease-onset to CSF sampling was 10 days (IQR: 5–25 days). The median white blood cell count in CSF was 80/mm^3^ (IQR: 19–220/mm^3^). The median CSF monocyte cell count was 36/mm^3^ (IQR: 10–127/mm^3^). During a follow-up period of 30 days, nine patients died.

**Figure 1.**
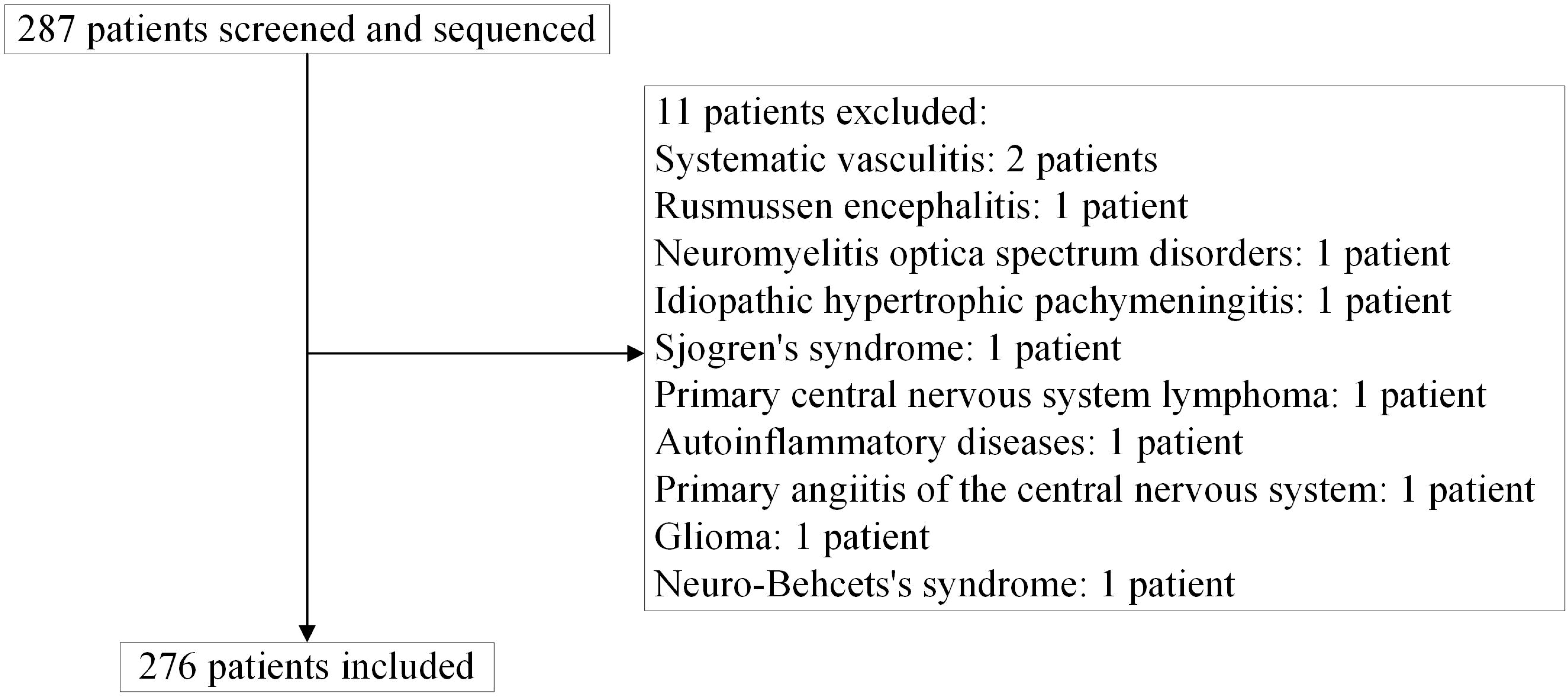
Flowchart of patient enrolment and exclusion.

### 3.2 Performance of mNGS for diagnosing CNS infections

276 CSF samples were tested by mNGS and conventional microbiological studies. 122 samples were positive by mNGS (110 true positive, 12 false positive), 126 were positive by conventional microbiological tests, and 114 total positive results were considered "Etiologic Pathogens" (Table 2). All mNGS results were obtained in less than 48 hours and 101 CSF samples were positive by mNGS before any conventional microbiological tests were positive.

**Table 2.** Performance of mNGS of CSF compared to conventional microbiological studies (gold standard) for the diagnosis of CNS infections

|  | TP* | FP | FN | Etiologic Pathogen |
| --- | --- | --- | --- | --- |
| **Bacteria** |  |  |  |  |
| *Mycobacterium tuberculosis* | 14 | 0 | 1 | 15 |
| *Listeria monocytogenes* | 8 | 0 | 0 | 8 |
| *Brucella* | 7 | 1 | 1 | 8 |
| *Streptococcus pneumoniae* | 5 | 0 | 0 | 5 |
| *Klebsiella pneumoniae* | 3 | 0 | 0 | 3 |
| *Streptococcus intermedius* | 2 | 0 | 0 | 2 |
| *Haemophilus influenzae* | 2 | 0 | 0 | 2 |
| *Vibrio vulnificus* | 1 | 0 | 0 | 1 |
| *Staphylococcus hominis* | 1 | 0 | 0 | 1 |
| *Escherichia coli* | 0 | 2 | 0 | 0 |
| *Enterococcus faecium* | 0 | 2 | 0 | 0 |
| *Acinetobacter baumannii* | 1 | 2 | 0 | 1 |
| *Stenotrophomonas maltophilia* | 1 | 1 | 0 | 1 |
| *Pseudomonas aeruginosa* | 0 | 4 | 1 | 1 |
| *Staphylococcus aureus* | 0 | 0 | 1 | 1 |
| *Staphylococcus haemolyticus* | 0 | 0 | 1 | 1 |
| **DNA Viruses** |  |  |  |  |
| Varicella-zoster virus | 17 | 0 | 4 | 21 |
| Herpes simplex virus 1 | 12 | 0 | 1 | 13 |
| Epstein-Barr virus | 12 | 0 | 3 | 6 |
| Cytomegalovirus | 4 | 0 | 0 | 2 |
| Herpes simplex virus 2 | 2 | 0 | 0 | 2 |
| *Suid herpesvirus 1* | 2 | 0 | 0 | 2 |
| BK polyomavirus | 1 | 0 | 0 | 0 |
| John Cunningham virus | 1 | 0 | 0 | 1 |
| **Fungi** |  |  |  |  |
| *Cryptococcus neoformans* | 7 | 0 | 3 | 10 |
| *Cryptococcus gattii* | 1 | 0 | 0 | 1 |
| **Parasites** |  |  |  |  |
| *Taenia solium* | 5 | 0 | 0 | 5 |
| *Angiostrongylus cantonensis* | 1 | 0 | 0 | 1 |
| **Totals** | **110** | **12** | **16** | **114** |

\* TP: true-positive; FP: false-positive; FN: false-negative; mNGS: metagenomic
next-generation sequencing; CSF: cerebrospinal fluid; CNS: central nervous system

Of the patients first diagnosed by mNGS, 16.3% of infections were caused by bacterial, 15.2% by viruses, 2.9% by fungi, and 2.2% by parasites (Fig. 2A).

**Figure 2.**
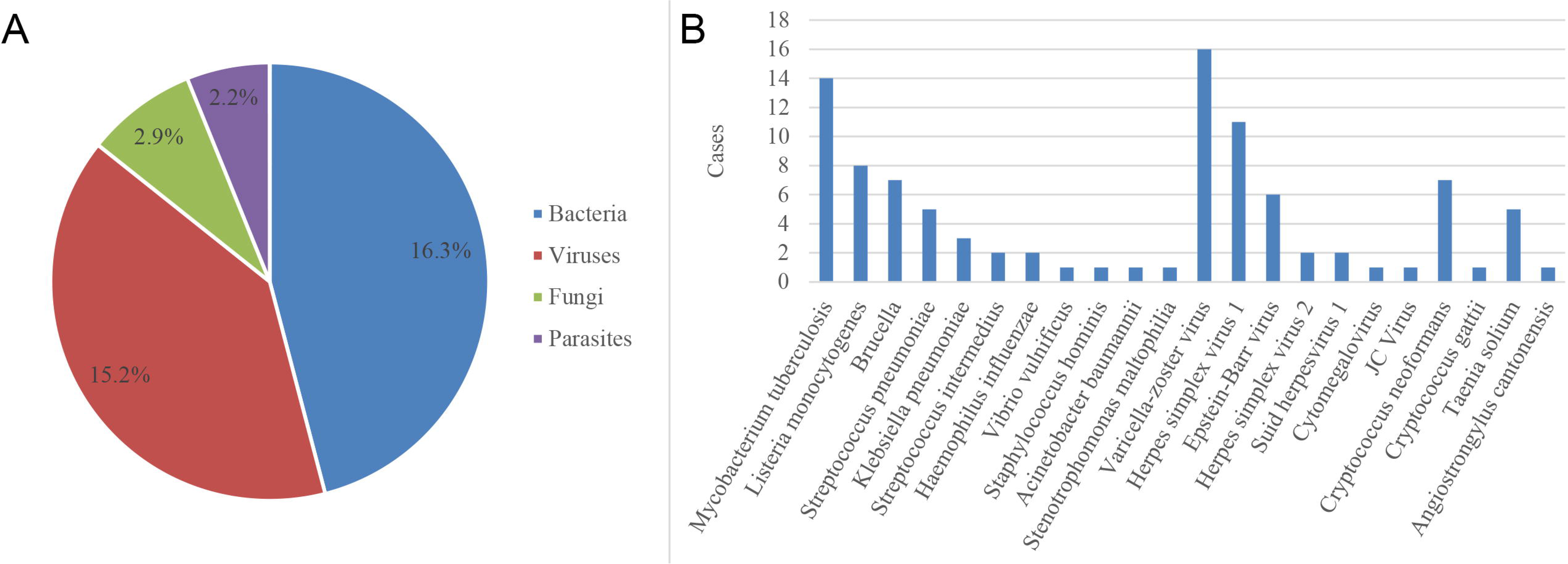
Distribution of causative pathogens in patients with suspected CNS infections initially detected by NGS of CSF. (A) Of the 36.6% patients first diagnosed with NGS of CSF, 16.3% were diagnosed with bacterial infections, 15.2% with viral infections, 2.9% with fungal infections, and 2.2% with parasitic infections. (B) NGS detected 11 bacterial species, the most common of which were *M. tuberculosis* (13.9%) and *L. monocytogenes* (7.9%), 7 viral species, the most common of which were VZV (16.8%) and HSV1 (11.9%), 2 fungal species, both of which were *Cryptococcus* (7.9%) species, and 2 parasitic species, the most common of which was *T. solium* (5.0%).

In total, NGS detected 11 bacterial species, of which *M. tuberculosis* (14 cases, 13.9%) and *L. monocytogenes* (7.9%) were the most common (Fig. 2B), 7 viral species (BK polyomavirus was not the etiologic pathogen), of which VZV (16.8%) and HSV1 (11.9%) were the most common, 2 fungal species, both of which were *Cryptococcus* (7.9%), and 2 parasitic species, of which *T. solium* (5.0%) was the most common. Nine co-infections with Epstein-Barr virus (EBV) (three with HSV1, two with *Brucella*, one with *Cryptococcus*, one with *S. haemolyticus*, one with *P. aeruginosa*, and one with *M. tuberculosis*), two co-infections with cytomegalovirus (CMV) (one with *M. tuberculosis*, and one with *Cryptococcus*), and one co-infection with BK polyomavirus (with HSV1) were detected. The EBV and BK polyomavirus did not appear to be consistent with the clinical manifestations in these two instances of co-infections.

### 3.3 False positive results of CSF mNGS

In the present study, false positives occurred in 12 (4.3%) patients and were primarily associated with bacterial infections (n=12; Table 2), including *E. coli*, *E. faecium*, *A. baumannii*, *S. maltophilia*, and *P. aeruginosa*, and a false positive for *Brucella* was also seen. Of note, the false-positive samples contained numerous other bacteria, that could be detected simultaneously by NGS. Using our proposed criteria, there were no false positives for viruses, fungi, or parasites. Although EBV was not the etiologic pathogen in most cases, it was present in the CSF of some patients. Additionally, there was some background contamination in most CSF samples (Supplementary Table 2) but these organisms did not meet the criteria for a positive result.

### 3.4 False negative results of CSF mNGS

In the present study, false negatives occurred in 16 (5.8%) patients (Table 2) and were associated with bacterial, viral and fungal infections. The false negative cases of bacterial infection were all treated with antibiotics prior to sequencing. In the false negative cases of viral infection, 1 or 2 SSRNs were detected in the samples but did not satisfy the proposed criteria for a positive mNGS result. If the criteria for a positive result was relaxed to a SSRN ≥ 1 (RPM ≥ 0.05), there were no false negative cases of HSV1 or VZV or false positive cases of HSV1, HSV2 or VZV. In this study, if we adopted the alternate criteria SSRN ≥ 1 (RPM ≥ 0.05) for viruses and *Mycobacterium tuberculosis*, there would be additional potential false positives, including 30 EBV, 7 CMV and 5 *Mycobacterium tuberculosis* infections. It should be pointed out that the possibility of *Mycobacterium tuberculosis* infection in the 5 cases cannot be ruled out based on the clinical and paraclinical manifestations, because the conventional microbiological methods might fail to detect the *Mycobacterium tuberculosis*. Of note, there were three false negative cases of *Cryptococcus* infection.

## 4. DISCUSSION

To the best of our knowledge, the present study is the first to assess the performance of mNGS for pathogen identification in a large prospective cohort of patients with suspected CNS infections. Specifically, our study compared results of mNGS of CSF to conventional microbiological studies and proposed new criteria for validating a mNGS result as positive for therapeutic decision-making. Our results suggest that NGS can provide a quicker and more accurate etiologic pathogen identification than conventional microbiological methods. However, patients in the present study were only eligible to be assessed by mNGS if conventional microbiological studies, e.g. routine bacterial stains and cultures, India ink preparation, targeted PCR tests, serological tests, failed to identify an etiologic cause within 3 days. Thus, the application of CSF mNGS in the clinical setting of this study could be regarded as a quasi-first line method for diagnosing CNS infectious diseases.

mNGS is a high-throughput sequencing technique without the requirement of prior information, allowing detection of unsuspected or novel organisms. Importantly, mNGS can detect unsuspected pathogens that clinicians may fail to consider because of atypical clinical manifestations. Many cases of neurological infections have been unexpectedly diagnosed by mNGS of CSF^11,22,29,30^ similar to the present study for the cases of *L. monocytogenes*, *Brucella* and *T. solium*^12–14^. In addition, as demonstrated in previous studies^10,20,21^ and in the present study for the case of encephalitis caused by *Suid herpesvirus 1* ^31^, mNGS of CSF has the ability to identify novel aetiologies of CNS infections. Furthermore, NGS can detect unexpected co-infections that may guide appropriate targeted treatment. For example, we detected co-infections of CMV and *Cryptococcus*. In routine clinical practice, if conventional microbiological methods detect *Cryptococcus*, then no further tests for other microorganisms other than HIV are usually performed. Finally, mNGS of CSF may be an appropriate tool for ruling out a broad spectrum of potential CNS infectious diseases prior to concluding a final diagnosis of autoimmune diseases, such as autoantibody-negative autoimmune encephalitis.

Contamination of samples during specimen collection and/or processing is a major challenge when interpreting mNGS results. To reduce the potential influence of contamination, we defined strict criteria for positive mNGS results. The various types of contamination observed in the present study could be divided into two groups: (1) microorganisms commonly associated with background contamination that did not meet the criteria for a positive result (Supplementary Table 2) and (2) false positive detections that fulfilled our criteria for a positive mNGS result but were not consistent with the patients’ clinical presentation and features. The contaminations derived primarily from the following sources: (1) laboratory practices (Parvovirus NIH CQV is a contaminant from silica column-based nucleic acid extraction kits)^32^; (2) reagents (*Bradyrhizobium*, *Burkholderia* and *Ralstonia* are common contaminants used in industrial ultrapure water systems)^33,34^; (3) environment (*E. coli*, *P. aeruginosa*, *E. faecium* and Torque teno virus are widespread pathogens in hospital environment)^35,36^; (4) skin or other body flora (*P. acnes*, *M. globose*, *E. coli* and *S. epidermidis* are widely associated with the human skin flora)^37,38^. False positive results are very likely to misguide treatment, and therefore, clinicians should be cautious when interpreting positive mNGS detection of extracellular bacteria or fungi that are widespread in hospital environments, especially when many species of bacteria are detected in a single NGS test. On the other hand, positive mNGS detection of viruses and parasites are not likely to be false positives.

Negative mNGS results do not necessarily exclude an infectious cause for the patient’s illness. In the present study, false negative mNGS results occurred in 5.8% of cases and included bacterial, viral, and fungal infections. The prior use of antibiotics can affect the detection rate of bacteria. Low SSRN values (1 or 2 reads) can be seen in false negative results for virus detection indicating that other microbiological tests should be conducted to confirm a diagnosis when the SSRN is low for viruses. False negative cases have been reported for *Cryptococcus* due to fungal counts below the lower limit of detection for nucleic acid amplification tests^39,40^. The criteria of SSRN ≥ 3 rather than SSRN ≥ 1 might introduce false-negative cases for *Mycobacterium tuberculosis* infections. As a screening method, the pathogens detected by mNGS might provide clinical clues for further investigation even if the results do not fulfil the specified criteria for a positive result.

Our results indicate that mNGS of CSF is a very useful test for the diagnostic evaluation of patients with suspected CNS infections. mNGS has already become a first-line laboratory method in the response to emerging infectious diseases and outbreaks of infectious diseases^41,42^. Moreover, increasing evidence provides a rationale for using mNGS as a first-line diagnostic test for chronic and recurring encephalitis and as a second-line test for acute encephalitis^23^. In the present study, more than one-third of patients were first diagnosed by mNGS of CSF within 48 hrs, indicating that this technique can be extremely useful for rapid clinical decision-making. mNGS of CSF should be considered as a first-line test for acute CNS infections when (1) a patient is critically ill and requires prompt and precise therapy, (2) the clinical manifestations are non-specific and numerous target-specific tests are simultaneously applied to identify an infectious cause, 3) a broad spectrum of potential pathogens needs to be ruled out to diagnose a suspected autoimmune encephalitis, and (4) rare or novel pathogens are suspected for which standard target-specific tests are not available.

There are several limitations regarding the current use of mNGS of CSF in routine clinical settings. First, mNGS is not available in many hospitals and the cost is usually much higher than target-specific tests. Second, the detection of DNA of a certain pathogen does not necessarily prove that it is responsible for the patient’s clinical presentation and features^23,27^. In the present study, some patients were positive for EBV but had other more likely aetiologies of their infections. EBV DNA has often been identified together with other microorganisms in CSF^43^. Finally, mNGS cannot detect microorganisms that are not included in microbial genome databases.

Our study has several limitations. Reverse transcription was not performed to construct a DNA library and therefore, RNA viruses could not be detected by mNGS. Next-generation RNA sequencing should be performed in future prospective studies. The number of CNS infections caused by each pathogen were not sufficient to assess the sensitivity and specificity of mNGS for individual pathogens. Finally, we may have underestimated the sensitivity of mNGS of CSF for diagnosing CNS infections by employing strict criteria for a positive mNGS result and using conventional microbiological methods as the gold standard.

## List of Abbreviations

mNGS: metagenomic next-generation sequencing
CSF: cerebrospinal fluid
CNS: central nervous system
PCR: polymerase chain reaction
HSV: herpes simplex virus
VZV: varicella zoster virus
NTC: ‘no-template’ controls
RPM: reads per million
EBV: Epstein-Barr virus
CMV: cytomegalovirus
SSRN: species-specific read number

## Declarations

### Consent for publication

Not applicable.

### Availability of data and materials

The datasets used and/or analysed during the current study are available from the corresponding author on reasonable request.

### Author’s contributions

HG and SF contributed to the study conception design. All the authors participated in the discussion of the study design. SF, XW, YF, JS, YZ, WZ, XQ, CW, LL, LQ, SW, HJ, HL, WY, WZ, XL, SL, YW, YP, HR, YZ, JW, BP, LC, WL, and HG enrolled patients and collected the clinical data. HW, YL and WC performed NGS, bioinformatics analysis, and PCR. SF, HG, XW, and WL analyzed the data. SF wrote the first draft of the manuscript after discussions with HG, XW and WL. HW wrote portions of the Methods section. HG, JHC, TG and WC contributed to manuscript revision. All authors have read and approved the submitted version.

## Acknowledgments

The authors thank the patients for participating in this study.

