## Supplementary Table 1 for "Metagenomic Next-generation Sequencing of Cerebrospinal Fluid for the Diagnosis of Central Nervous System Infections: A Multicentre Prospective Study"

Supplementary Table 1. Routine diagnostic examinations in the present study.

**Peripheral blood**

Routine blood cultures

HIV serology, Treponemal testing

ANA

Anti-NMDAR, anti-Caspr2, anti-AMPAR 1, anti-AMPAR 2, anti-LGI1, and anti-GABA_B_-R antibodies; anti-CV2/CRMP5, anti-PNMA2, anti-Ri, anti-Yo, anti-Hu, and anti-Amphiphysin antibodies

**CSF**

Opening pressure, WBC count with differential, RBC count, protein, chloride, glucose

Gram stain and bacterial culture, India Ink staining, acid-fast staining

Anti-NMDAR, anti-Caspr2, anti-AMPAR 1, anti-AMPAR 2, anti-LGI1, and anti-GABA_B_-R antibodies; anti-CV2/CRMP5, anti-PNMA2, anti-Ri, anti-Yo, anti-Hu, and anti-Amphiphysin antibodies

**Imaging**

Neuroimaging (MRI preferred to CT, if available)

Chest imaging (Chest x-ray and/or CT)

**EEG**

**Conditional examinations**

Epidemic season: Japanese encephalitis virus IgM testing

Epidemic area: brucella testing

TB testing when necessary

AMPAR: α-amino-3-hydroxy-5-methyl-4-isoxazolepropionic acid receptor; HIV: human immunodeficiency virus; ANA: antinuclear antibody; NMDAR: N-methyl-D-aspartate receptor; Caspr2: contactin-associated protein-like 2; LGI1: leucine-rich, glioma inactivated 1; GABA: gamma-Aminobutyric acid; CRMP5: collapsin response-mediator protein-5; PNMA2: paraneoplastic antigen Ma2; MRI: magnetic resonance imaging; CT: computerized tomography; CSF: cerebrospinal fluid; EEG: electroencephalogram; WBC: white blood cell.
