## Supplementary Table 2 for "Metagenomic Next-generation Sequencing of Cerebrospinal Fluid for the Diagnosis of Central Nervous System Infections: A Multicentre Prospective Study"

Supplementary Table 2. Common background contaminations of CSF mNGS

**Bacteria**

*Propionibacterium acnes*

*Propionibacterium avidum*

*Acidovorax KKS102*

*Methylobacterium radiotolerans*

*Bradyrhizobium japonicum*

*Bradyrhizobium S23321*

*Bradyrhizobium BTAi1*

*Staphylococcus epidermidis*

*Staphylococcus haemolyticus*

*Staphylococcus capitis*

*Ralstonia solanacearum*

*Micrococcus luteus*

*Burkholderia multivorans*

*Burkholderia ambifaria*

*Burkholderia cepacia*

*Burkholderia 383*

*Sphingobium japonicum*

*Sphingopyxis alaskensis*

*Brevundimonas subvibrioides*

*Asticcacaulis excentricus*

*Cupriavidus metallidurans*

*Comamonas testosterone*

*Acinetobacter lwoffii*

**Fungi**

*Malassezia globose*

*Saccharomyces bayanus*

**Parasites**

*Trichinella papuae*

*Trichinella zimbabwensis*

*Echinococcus granulosus*

**Viruses**

*Torque teno virus*

*Parvovirus NIH CQV*

NGS: next-generation sequencing; CSF: cerebrospinal fluid.
