## Supplementary Table 3 for "Metagenomic Next-generation Sequencing of Cerebrospinal Fluid for the Diagnosis of Central Nervous System Infections: A Multicentre Prospective Study"

Supplementary Table 3. Species-specific reads numbers of CSF NGS for the true-positive, false-positive and false-negative cases.

|  | **SSRNs for TP,**  **range/no.** | **SSRNs for FP,**  **range/no.** | **SSRNs for FN,**  **range/no.** |
| --- | --- | --- | --- |
| **Bacteria** | | | |
| *Mycobacterium tuberculosis* | 3-396 | - | 0 |
| *Listeria monocytogenes* | 307-1591 | - | - |
| *Brucella* | 5-1417 | 5 | 2 |
| *Streptococcus pneumoniae* | 788-688437 | - | - |
| *Klebsiella pneumoniae* | 173-1011 | - | - |
| *Streptococcus intermedius* | 138-591 | - | - |
| *Haemophilus influenzae* | 345-19239 | - | - |
| *Vibrio vulnificus* | 1047 | - | - |
| *Staphylococcus hominis* | 34 | - | - |
| *Escherichia coli* | - | 420-1089 | - |
| *Enterococcus faecium* | - | 249-456 | - |
| *Acinetobacter baumannii* | 5481 | 322-103660 | - |
| *Stenotrophomonas maltophilia* | 19848 | 231 | - |
| *Pseudomonas aeruginosa* | - | 264-1591 | 2 |
| *Staphylococcus aureus* | - | - | 1 |
| *Staphylococcus haemolyticus* | - | - | 1 |
| **Viruses** | | | |
| Varicella-zoster virus | 3-51731 | - | 1-2 |
| Herpes simplex virus 1 | 4-31402 | - | 2 |
| Epstein-Barr virus | 3-38 | - | 1 |
| *Cytomegalovirus* | 3-10 | - | - |
| Herpes simplex virus 2 | 10-332 | - | - |
| *Suid herpesvirus 1* | 20-242 | - | - |
| *BK polyomavirus* | 4 | - | - |
| *JC Virus* | 4 | - | - |
| **Fungi** | | | |
| *Cryptococcus neoformans* | 33-226572 | - | 0 |
| *Cryptococcus gattii* | 68261 | - | - |
| **Parasites** | | | |
| *Taenia solium* | 108-66508 | - | - |
| *Angiostrongylus cantonensis* | 718 | - | - |

TP: true positive; FP: false positive; FN: false negative; no.: number; NGS: next-generation sequencing; CSF: cerebrospinal fluid, SSRN: Species-specific read number
